# Tumor-specific Kinase Motif Enrichment Analysis Identifies Personalized Therapeutic Cancer Targets

**DOI:** 10.64898/2026.07.31.742097

**Authors:** Tracey Pu, Brian A. Joughin, Priyanka P. Desai, Steven D. Forsythe, Ronald Holewinski, Kirsten Remmert, Billel Gasmi, Kendra Coleman, Timothy Leach, Allan M. Johansen, Nicole Russell, Lichun Ma, Ashley Rainey, Amber Leila Sarvestani, Emily Smith, Surajit Sinha, Sunanda Mukherjee, Kenneth Luberice, Sophia Xiao, Carolina M. Larrain, Alyssa V. Eade, Lindsay R. Friedman, Jason Ho, Jeremy L. Davis, Andrew M. Blakely, David E. Kleiner, Samira M. Sadowski, Thorkell Andresson, Jaydira Del Rivero, Michael B. Yaffe, Jonathan M. Hernandez

**Affiliations:** Surgical Oncology Program, Center for Cancer Research, National Cancer Institute, National Institutes of Health, Bethesda, MD, USA; Koch Institute for Integrative Cancer Research, Departments of Biological Engineering and Biology, Massachusetts Institute of Technology, Cambridge, MA, USA; Neuroendocrine Cancer Therapy Section, Center for Cancer Research, National Cancer Institute, National Institutes of Health, Bethesda, MD, USA; Protein Characterization Laboratory, Frederick National Laboratory for Cancer Research, Leidos Biomedical Research, Frederick, Maryland, USA; Surgery Branch, National Cancer Institute, National Institutes of Health, Bethesda, MD, USA; Wake Forest School of Medicine, Medical Center, Wake Forest Institute for Regenerative Medicine, Winston-Salem, NC, USA; Department of Oncology, School of Medicine, Karmanos Cancer Institute, Wayne State University, Detroit, MI, USA; Cancer Data Science Laboratory, Center for Cancer Research, National Cancer Institute, Bethesda, MD, USA; Liver Cancer Program, Center for Cancer Research, National Cancer Institute, Bethesda, MD, USA; Laboratory of Pathology, National Cancer Institute, NIH, Bethesda, Maryland, USA; Developmental Therapeutics Branch, Rare Tumor Initiative, Center for Cancer Research, National Cancer Institute, National Institutes of Health, Bethesda, MD, USA; Divisions of Surgical Oncology, Acute Care Surgery, Trauma, and Surgical Critical Care, Beth Israel Deaconess Medical Center, Harvard Medical School, Boston, MA, USA

## Abstract

Gastroenteropancreatic neuroendocrine tumors (GEP-NETs) are an uncommon and poorly understood malignancy with low mutational burden, lacking well-defined oncogenic drivers. GEP-NET mortality frequently results from extensive hepatic metastases. Accordingly, we interrogated phosphoproteomic data from GEP-NET liver metastases and patient-matched uninvolved liver to identify tumor-specific signaling and targetable tumor vulnerabilities using Kinase Motif Enrichment Analysis (KMEA), a new tool leveraging the recent Kinase Library compendium of the substrate motif specificity for nearly the entire human kinome. KMEA identified patient tumor-specific upregulation of mTOR or casein kinase 2 (CK2) activity that would be undiscoverable by standard personalized genomic and transcriptomic approaches. Striking concordance was observed between KMEA predictions for specific tumors, and their sensitivity to inhibitors of mTOR or CK2 using patient tumor-derived organoids. These findings reveal potential clinically-actionable protein kinases hyperactivated in GEP-NETs, and more broadly indicate a general method for personalized cancer treatment using phosphoproteomics and KMEA-derived kinase activity signatures.

## INTRODUCTION

Tumor characterization technologies have enabled oncology to rapidly move towards personalized medicine [1]. Insights from next-generation sequencing (NGS) have revealed conserved mutations in driver oncogenes such as *KRAS, BRAF, EGFR, ALK,* and *PIK3CA*, which are now targetable with an expanding armamentarium of inhibitors [2, 3]. Consequently, solid tumor oncology has incorporated molecular targets in combination with histological characterization to select patients for therapies [4–6]. For example, significant and durable responses to c-Kit tyrosine kinase inhibitors have been observed for patients with gastrointestinal stromal tumors harboring specific exon mutations [7, 8]. Despite rapid adoption of NGS and comprehensive genomic profiling, multiple recent real-world studies and systematic reviews indicate that sequencing alone has not produced consistent, population-level improvements in cancer outcomes [9–11]. Meta-analyses and pooled real-world data show that only a minority of patients (often in the range of ∼10–30%) are ultimately treated with genomics-matched therapies, and reported survival gains are heterogeneous, even when confined to well-validated predictive biomarkers (e.g. *EGFR, ALK, HER2, NTRK*) [10, 11].

Several factors may help explain the discrepancy between genomic insight and broad clinical impact. These include the modest prevalence of truly actionable, predictive genomic alterations in driver oncogenes found in unselected advanced-cancer populations [12] such as lung adenocarcinoma in never-smokers without *EGFR* or *ALK* mutations [13] or small bowel neuroendocrine tumors [14]. Additionally, identification of a targetable mutation is often taken as synonymous with reliance on that target/pathway for tumor cells survival/viability, although confirmatory measurements of expression and pathway activity are typically not performed [15]. For example, response to BRAF/MEK inhibitors for patients with *BRAF*^V600E^-mutated melanoma is 60-70% [16], while that response rate drops to 12% for patients with *BRAF*^V600E^-mutated colorectal cancer [17]. Therefore, relying on mutational data alone to infer potential drivers cannot guarantee success. Other tumor types, including high grade serous ovarian cancer [18], endometrial cancer [19] and prostate cancer [20], lack oncogenic drivers and instead predominantly contain mutations in tumor suppressor genes such as *TP53, ATM*, or *PTEN* for which effective therapeutic approaches that restore wild-type function are lacking.

An alternative approach to identify and target critical tumor dependencies involves the use of proteomics and phosphoproteomics to directly measure the signaling state of the tumor [21–27]. Proteomic, rather than purely genomic and transcriptomic signatures, are not the current standard of care in the clinic, but have been used in specialized research trials, and in advanced molecular tumor boards, rather than as a routine frontline diagnostic tool for all cancer patients [11, 28–31]. Clinical implementation of mass spectrometry-based proteomic technologies remains a difficult challenge owing to technical complexities, hardware costs, and the stochastic nature of phosphopeptide identification in discovery-mode mass spectrometry, although significant advances incorporating this technology into clinical pipelines are being made [32]. Even with this progress, a significant hurdle remains in converting lists of quantified phosphopeptides from clinical samples into potentially actionable patient treatments targeting tumor-specific kinase dysregulation [33].

A tumor-specific list of phosphopeptides, rather than a genomic signature, could have the potential for rapid clinical translation, if the relationships between the measured phosphopeptides and the protein kinases responsible for their phosphorylation were well-understood. Although the human proteome encodes over 500 protein kinases, there are very few known substrates for most of them [33–35]. Conversely, modern mass spectrometry techniques have identified over 200,000 human phosphorylation sites, but the specific protein kinase responsible for phosphorylation is unknown for more than 90% of the sites [36, 37]. These gaps in knowledge represent a massive roadblock in translating discovery-mode mass spectrometry into therapeutically actionable targeting of kinase dysregulation in cancer.

Here we present Kinase Motif Enrichment Analysis (KMEA), a software tool that leverages our recently reported compendium of protein kinase phosphorylation motifs for the serine/threonine kinome [38] to interrogate tumor phosphoproteomic data. We focus on characterizing hepatic metastases from sporadic gastrointestinal and pancreatic neuroendocrine tumors (GEP-NETs) given our experience modeling this cancer [39] for personalized medicine approaches coupled with our clinical expertise with this patient population [40–46]. Importantly, GEP-NETs arise secondary to non-genomic alterations in signaling pathways that control cell proliferation, survival and invasion as a consequence of epigenetic changes, differences in post-transcriptional regulation, and/or pathway re-wiring to create novel feedback loops and/or dependencies [47–50] making them ideal for personalized signal transduction interference strategies. We show that KMEA-based interrogation of GEP-NET phosphoproteomic data, combined with patient-derived tumor avatar validation, successfully identified tumor-specific signaling dependencies that were not evident from genome-based interrogation methods.

## RESULTS

### GEP-NETs demonstrate genetic heterogeneity with limited actionable targets via next-generation sequencing (NGS)

Comprehensive genomic profiling consistent with current care guidelines was performed on liver metastases from 15 patients with small bowel (SBNET) or pancreatic (PNET) neuroendocrine tumors who underwent hepatic metastatectomy. Figures 1A and 1B show mutations identified in SBNET and PNET patients, respectively, using a comprehensive TruSight Oncology 500 NGS assay that utilizes a hybrid-capture approach to concurrently assess DNA and RNA for the detection of small variants, fusions, and copy number alterations across 523 cancer-related genes, while providing quantitative assessment of tumor mutational burden and microsatellite instability [51]. Only three somatic mutations were observed in SBNETs: *CDKN1B* (2/8 patients) and *DAXX* (1/8) (Fig. 1A). In PNETs, more than half the tumors showed evidence of somatic MEN1 mutations, while other common mutations included *DAXX, SETD2*, and *TP53*, consistent with previous reports [52–54]. Only three patients were matched to an FDA-approved therapy on the basis of mutational profile, which was the mTOR inhibitor everolimus [55] (Fig. 1C). For the remaining 12 (80%) patients, no therapy was suggested on the basis of genomic analysis. Details regarding the use of samples and data from each patient throughout this manuscript are given in Supplemental Figure 1.

**Figure 1.**
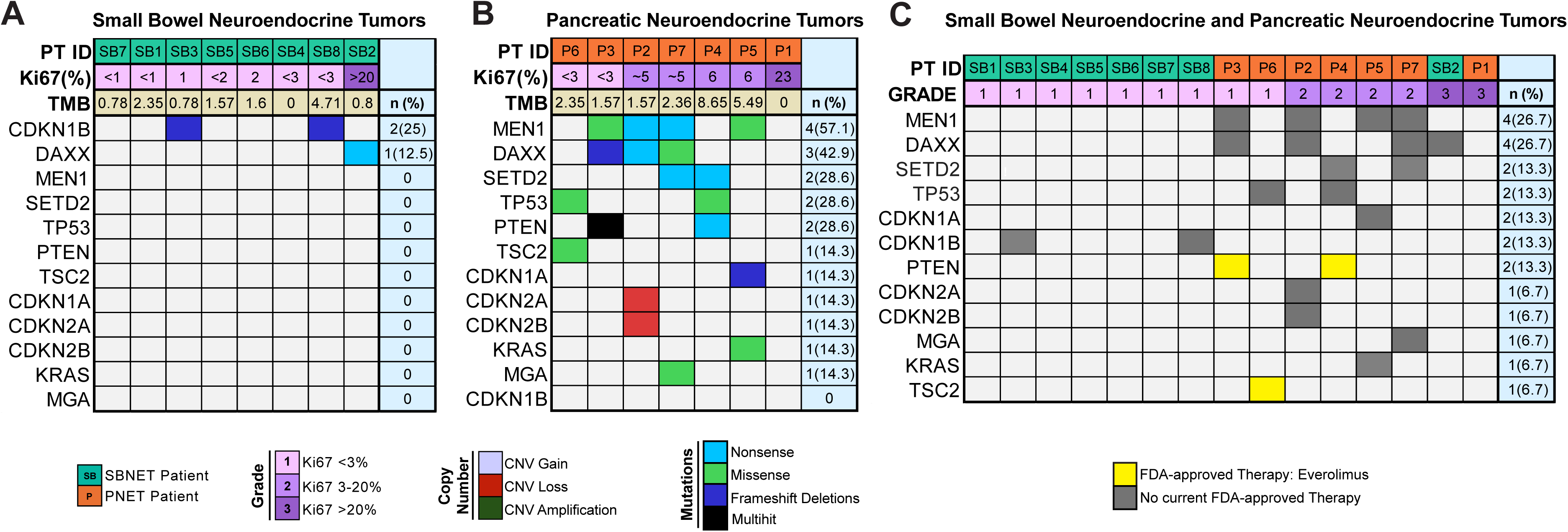
GEP-NETs demonstrate genetic heterogeneity with limited actionable targets via next generation sequencing (NGS). Oncoplot showing pathogenic mutations from **A.** SBNET (*n=8*) and **B.** PNET (*n=7*) liver metastases with respective percentages of Ki67 and tumor mutational burden (TMB). **C**. Oncoplot highlighting FDA-approved (yellow) drug availability according to pathogenic mutations (*n=15* patients).

### KMEA Identifies Upregulated Signaling Pathways and Potentially Translatable Targets in GEP-NETs Liver Metastases

In an effort to non-genomically identify kinases whose dysregulation may be important to tumor growth and survival, SBNET and PNET hepatic metastases and patient-matched uninvolved liver were snap-frozen in the operating room for proteomic and phosphoproteomic analysis by mass spectrometry (Schematized in Fig. 2A). Using tandem mass tag (TMT) mass spectroscopy to compare metastatic tumor tissue to uninvolved liver, we identified and quantified with high confidence a total of 2300 phosphorylated, and 16609 non-phosphorylated peptides across 10 metastases from 8 patients with SBNETs, as well as 2517 phosphopeptides and 17378 non-phosphorylated peptides across 9 metastases from 7 patients with PNETs (see Methods, Supplemental Table 1). Bulk analyses of this proteomic data compared the mean level of phosphopeptides across all patient metastases to the mean values across all liver control tissue samples. While the average non-phosphorylated peptide was downregulated 0.85- fold in SBNET liver metastases or upregulated 1.10-fold in PNET liver metastases vs. patient-matched uninvolved liver (Supplemental Fig. 2), a much higher and more consistent upregulation of phosphopeptides was observed: 4.09-fold in SBNETs and 2.83-fold in PNET liver metastases vs. patient-matched uninvolved liver (Fig. 2B).

**Figure 2.**
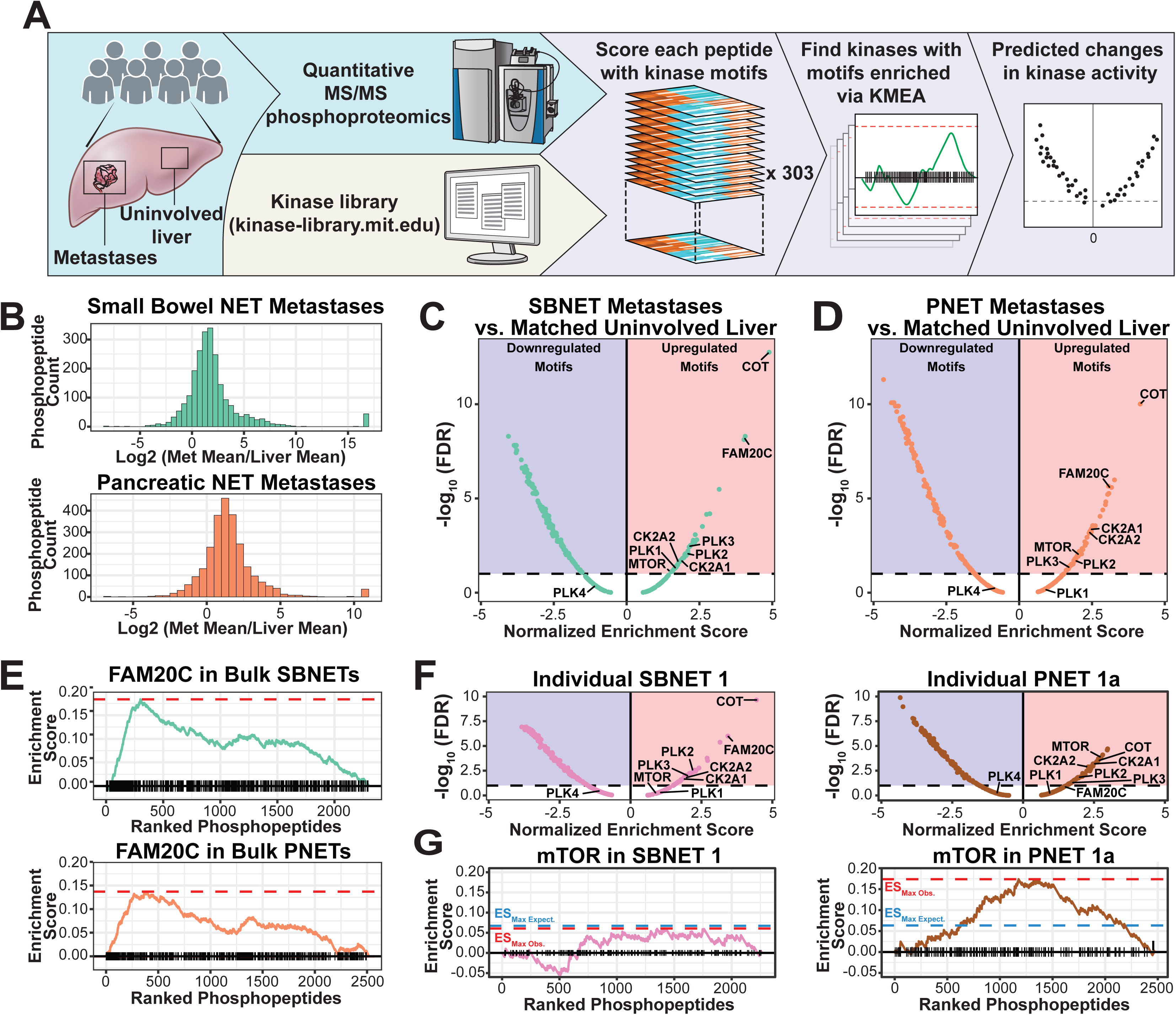
KMEA reveals potentially translatable targets in GEP-NET liver metastases. **A.** Schematic of KMEA analytical methods showing snap-frozen patient-matched uninvolved liver and NET liver metastases subjected to quantitative phosphoproteomic tandem mass spectrometry. Resultant phosphopeptide sequences are analyzed using the Kinase Library to associate each phosphopeptide with kinases that may have phosphorylated it. KMEA then identifies kinases whose substrate motifs are most associated with up- or downregulated phosphopeptides and thus, generates hypotheses regarding dysregulated kinases. **B.** Histograms of log_2_ fold-change in phosphopeptides in lesion relative to liver control for bulk SBNET (top) and PNET (bottom) liver metastases. **C,D.** Volcano plot indicating normalized enrichment score (NES) and -log_10_(FDR) for 303 human serine/threonine kinases in bulk SBNET (C) and PNET (D) liver metastases vs patient-matched uninvolved liver (control). Dashed line indicates an FDR of 0.1. Selected kinases are indicated. **E.** Running enrichment plots for the kinase motif set for the kinase FAM20C in bulk SBNET (top) and bulk PNET (bottom) vs. patient-matched uninvolved liver. Hash marks on the x-axis indicate phosphopeptides that may be phosphorylated by FAM20C in a list of phosphopeptides sorted by log_2_ fold-change. Red dashed lines indicate the maximum value of running enrichment score (ES) for the kinase motif set in the data. **F.** As in (C,D) but for a single SBNET (left) and a single PNET (right). **G.** As in (E) but for the mTOR kinase motif set in a single SBNET (left) and a single PNET (right). Blue dashed lines indicated the expected ES value from a distribution of permuted negative controls.

In order to explore kinase dysregulation in these two patient populations, the sequence of each phosphopeptide in the phosphoproteomic dataset was matched to its potential upstream kinases based on optimal amino acid substrate motifs determined for 303 serine/threonine kinases in the Kinase Library (https://kinase-library.mit.edu) [38, 56]. All phosphopeptides containing at least one phosphorylation site scoring in the 95^th^ percentile or better [38] for a given kinase were compiled into a “kinase motif set” for that kinase, and then data was analyzed using a computational framework comparable to that used for Gene Set Enrichment Analysis (GSEA) [57], which we term Kinase Motif Enrichment Analysis (see Methods). The output of KMEA is, for each kinase, an enrichment score (ES) calculated as the maximum distance from zero of a running enrichment score along a list of phosphopeptides, rank-ordered from most upregulated to most downregulated. A step up in running enrichment for a kinase is taken when a peptide is a putative kinase substrate, and a step down is taken when it is not. This maximum enrichment is transformed to a normalized enrichment score (NES) describing the degree to which a set of the phosphopeptides associated with a kinase is over- represented to one end or the other of a list of all phosphopeptides ranked by fold-change, relative to an identically sized random set of phosphopeptides. Positive NES values indicate a kinase’s substrate specificity motif is overrepresented among more upregulated phosphopeptides, whereas a negative NES indicates overrepresentation among the most downregulated phosphopeptides. A *p*-value is also calculated indicating the estimated proportion of random sets of the same size that would receive a given NES value, as well as a false discovery rate (FDR) value calculated by the method of Benjamini & Hochberg [58], acting as a multiple hypothesis correction across the 303 kinases. These data are used to generate volcano plots that nominate putatively dysregulated kinases based on the magnitude (NES) and significance (FDR) of their up- and downregulated motifs in the phosphoproteomic data (Fig. 2C-D).

Strikingly, the substrate specificity motifs for the kinases COT and FAM20C were among the three most significantly associated with phosphopeptide upregulation in both SBNETs (Fig. 2C) and PNETs (Fig. 2D). COT is a MAPKKK previously identified as a driver of resistance to RAS inhibition in melanoma cell lines [59]. FAM20C is a kinase active in the Golgi apparatus that is responsible for phosphorylating secreted proteins [60], an interesting confirmatory result consistent with cells of neuroendocrine origin having high secretory function [61]. Notably, potential substrates of FAM20C are highly enriched in roughly the top 20% of upregulated phosphopeptides in SBNETs, as can be seen by the sharp rise of a running enrichment plot among the 400-500 most upregulated phosphorylated peptides (Fig. 2E).

Although clinical inhibitors do not exist for COT or FAM20C, the serine/threonine kinase mTOR is inhibited by everolimus, a small molecule currently in clinical use as an FDA-approved standard-of-care treatment for NETs [62]. KMEA identified the mTOR substrate specificity motif as being significantly associated with upregulated peptides in both SBNETs (NES=1.58, FDR=0.07, Fig. 2C) and PNETs (NES=2.05, FDR=0.01, Fig. 2D) at an FDR cutoff of less than 10%. The higher degree of mTOR activity that we observe in PNETs as compared to SBNETs is consistent with the published observation that foregut-derived NETs have higher mTOR expression and activity than midgut-derived NETs [47, 63].

We next explored whether there was interpatient heterogeneity in mTOR activity by applying KMEA to 19 individual patient metastases with patient-matched uninvolved liver as a control. Indeed, we found that 6/9 PNET metastases had statistically significant (FDR < 10%) upregulation of phosphopeptides predicted to be mTOR substrates, and by contrast the same was true in only 2/10 SBNET metastases. Examples of two individual analyses, one of an SBNET with no signature of mTOR dysregulation (NES=0.90, FDR=0.62) and one of a PNET with putatively upregulated mTOR (NES=1.53, FDR=0.09) are shown in Figure 2F. The maximum running enrichment score for the SBNET (0.061) is less than that expected by chance, whereas maximum running enrichment score for the PNET (0.174) is over 250% of what would be expected by chance (Fig. 2G). Bulk SBNET, PNET, and individual patient phosphopeptides ratios are given in Supplemental Table 2. Complete KMEA results for bulk SBNET, bulk PNET, and each individual metastasis are given in Supplemental Table 3.

### Patient-derived organoids validate KMEA-nominated personalized targets in GEP-NETs Liver Metastases

KMEA analysis indicated statistically significant upregulation of mTOR substrate phosphorylation motifs in some, but not all, SBNET and PNET tumors (7/19), indicating the potential for KMEA to inform phosphoproteomically-driven personalized therapy. We therefore sought to functionally interrogate the interpatient differences in mTOR dependency that were suggested by patient-specific KMEA analysis. Preclinical NET models, however, are notoriously difficult to generate, and NET cell lines are extremely limited in number relative to other cancer types [46]. Given our expertise with NET organoid culture [39] and the personalized nature of KMEA predictions, we used patient-derived organoid (PDO) models to explore whether KMEA-predicted upregulation of mTOR activity correlated with sensitivity to everolimus (schematized in Fig. 3A).

**Figure 3.**
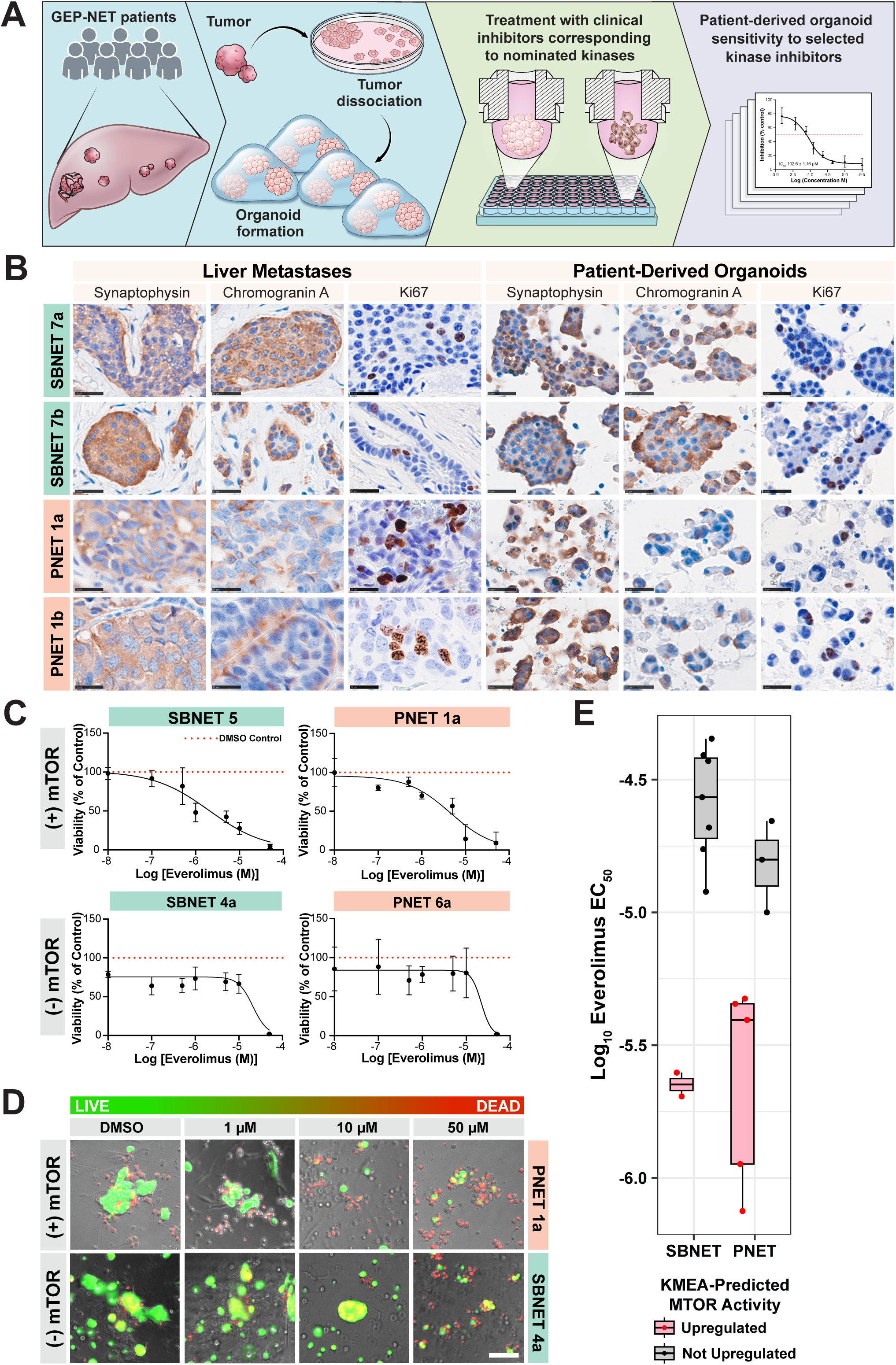
KMEA identifies mTOR therapeutic vulnerability in a subset of GEP-NET Liver Metastases. **A**. Schematic representation showing procurement of GEP-NET patient-derived liver metastases and patient-matched uninvolved liver from the operating room. Tumors are dissociated into single cell suspension for organoid preparation. After patient-derived organoid (PDOs) formation, PDOs are treated with selected clinical inhibitors based on KMEA. **B**. Representative images of IHC characterization of synaptophysin, chromogranin A, and Ki67 in SBNET (top, *n=2*) and PNET (bottom, *n=2*) liver metastases (left) and PDOs (right). Scale bar, 25 μm **C.** Representative dose-response curves of everolimus treated SBNET (left) and PNET (right) PDOs stratified by mTOR prediction by KMEA and represented based on mTOR expression as mTOR positive (mTOR+; top, *n=2*) and mTOR negative (mTOR-; bottom, *n=2*). Red dotted line represents control**. D**. LIVE /DEAD cell imaging using calcein-AM (green) and ethidium homodimer (red) showing dose-dependent cytotoxicity response in everolimus-treated SBNET (top) and PNET (bottom) PDOs and represented as mTOR positive (mTOR+; top) and mTOR negative (mTOR-; bottom). Ex: 488 nm, Em: 515 nm - calcein-AM; Ex: 570 nm, Em: 602 nm - Ethidium homodimer; Scale bar, 100 μm. **E.** Box-and-whisker plots indicating the half maximal effective concentration (EC_50_) values from SBNET (n=9) and PNET (n=8) PDOs after treatment with everolimus. Boxes indicate the median, 25^th^ and 75^th^ percentile values and whiskers extend to a maximum of 1.5 times the interquartile range. Data are separated by tissue of tumor origin and by whether KMEA predicted upregulation of mTOR. A single PNET EC_50_ which could not be confidently fit was manually placed at 10μM, visually estimated as the lowest-bound value consistent with data.

In a variety of other tumor types, PDOs have been shown to predict patient responses to targeted therapy with high sensitivity and specificity through maintenance of 3D spatial architecture and approximation of tumor heterogeneity [64–67] and this has recently been extended to NETs [68, 69]. Accordingly, we successfully established PDO models for 17 out of the 19 tumor metastases evaluated by KMEA, using our previously published technique [39]. In brief, PDOs were cultured until the formation of large clusters of cells were detected, or until lack of proliferation exceeding 60 days was noted. Immunohistochemistry (IHC) markers, including chromogranin A, synaptophysin, and Ki67 were used to assess neuroendocrine phenotype and showed similar marker expression to matched tumors (Fig. 3B, Supplemental Fig. 3). Pharmacologic testing of the organoids using varying doses of everolimus was undertaken for all PDO models. Viability was quantitatively assessed using CellTiter Glo after 96 hours of treatment (Fig. 3C and Supplemental Fig. 4) and corroborated by visual assessment with imaging of live cells by calcein AM and dead cells by ethidium homodimer (Fig. 3D, green and red cells, respectively). Dose-response curves were fit to the Cell-Titer Glo data, revealing a systematic difference between PDOs generated from tumors where KMEA nominated mTOR as being upregulated, compared to PDOs generated from tumors without mTOR nomination (Fig. 3E). Specifically, the EC_50_ in PDOs generated from tumors where KMEA nominated mTOR ranged from 0.75μM to 4.74μM, while PDOs generated from tumors without mTOR nomination had EC_50_ in a non-overlapping range from 10μM to 45μM with the medians of these distributions differing by roughly one order of magnitude (Fig. 3E).

Given the ability of KMEA to predict patient-specific PDO mTOR activity, we next examined whether there were other druggable kinases with enhanced activity in NETs. We identified casein kinase 2 (CK2A1, CK2A2) as a kinase that showed greater upregulation than mTOR in both the bulk SBNET and bulk PNET phosphoproteomic analyses (see Fig. 2C-D), and as a target kinase with an inhibitor that has received orphan drug designation by the FDA [70]. CK2 is a constitutively expressed serine/threonine kinase that is known to stabilize signaling networks that support cell growth and survival, and is upregulated in many tumor types [71]. Interestingly, CK2 expression was consistently observed to be higher in hepatic GEP-NET metastases by IHC than in patient-matched uninvolved liver (Fig. 4A and Supplemental Fig. 5) and this expression was preserved in PDOs (Fig. 4B). Despite constitutive expression, regulation of CK2 activity is complex and incompletely understood, involving a combination of kinase- intrinsic and extrinsic factors [72]. Significant enrichment of phosphopeptides predicted to be CK2 substrates was seen in 5/9 SBNET liver metastases and 7/10 PNET liver metastases as compared to patient-matched uninvolved liver, despite high CK2 expression by IHC in all PNET liver metastases and most SBNET liver metastases. Importantly, we noted that CK2 was nominated by KMEA as being specifically activated in a subset of individual patients whose organoids were insensitive to the mTOR inhibitor everolimus, suggesting CK2 as a potential therapeutic target in this population (Fig. 4C).

**Figure 4.**
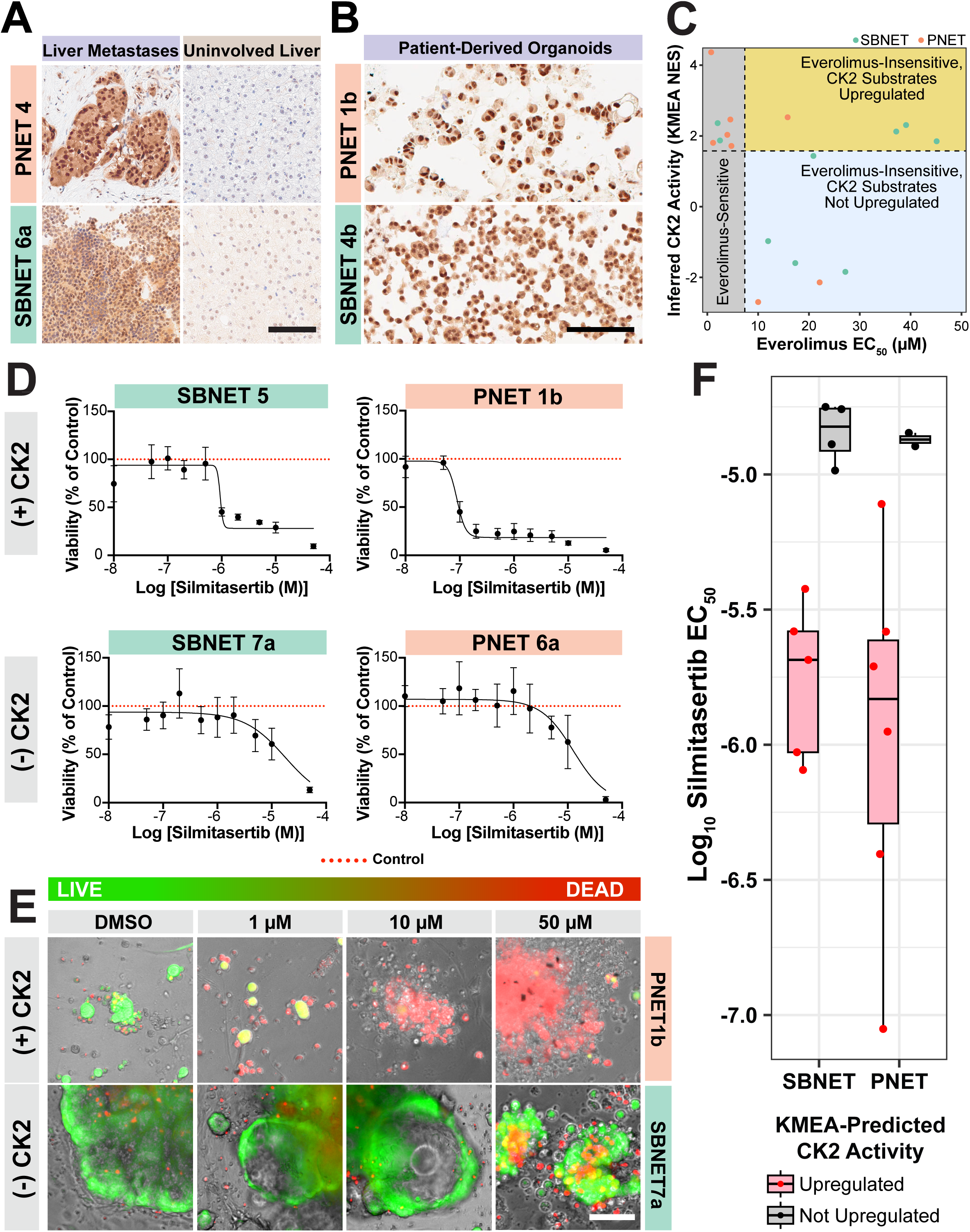
KMEA reveals casein kinase 2 as a novel therapeutic target in GEP-NET liver metastases. Representative images of IHC characterization of Casein Kinase 2 (CK2) staining in **A**. PNET (top) and SBNET (bottom) patient-derived liver metastases (left) and patient-matched uninvolved liver (right) and **B.** PNET (top) and SBNET (bottom) PDOs. Scale bar, 25 μm **C.** Scatter plot indicating everolimus EC_50_ of each PDO model and KMEA NES for CK2 in the corresponding liver metastases vs. patient-matched uninvolved liver as a control. Points are colored by tumor tissue of origin, and background coloring indicates regions where PDOs are relatively sensitive to everolimus, or insensitive to everolimus with or without a prediction of high CK2 activity. **D.** Representative dose-response curves of silmitasertib-treated SBNET (left) and PNET (right) PDOs stratified by CK2 prediction by KMEA and represented as CK2 positive (CK2+; top, *n=2*) and CK2 negative (CK2-; bottom, *n=2*). Red dotted line represents control. **E**. LIVE /DEAD cell imaging using calcein-AM (green) and ethidium homodimer (red) showing dose-dependent viability response in silmitasertib-treated PNET (top) and SBNET (bottom) PDOs and represented as CK2 positive (CK2+; top) and CK2 negative (CK2-; bottom). Ex: 488 nm, Em: 515 nm - Calcein-AM; Ex: 570 nm, Em: 602 nm - Ethidium homodimer; Scale bar, 100 μm. **F.** Box-and-whisker plots indicating the EC_50_ values from each SBNET (n=9) and PNET (n=8) PDOs after treatment with silmitasertib. Boxes indicate the median, 25^th^ and 75^th^ percentile values of the associated data, and whiskers extend to a maximum of 1.5 times the interquartile range. Data are separated by tissue of tumor origin and by whether KMEA predicted upregulation of CK2.

All SBNET and PNET PDOs were therefore treated with increasing doses of silmitasertib, an ATP-competitive small-molecule CK2A/B inhibitor with reported antiproliferative effects in a subset of cancer model cell lines [73]. Silmitasertib demonstrated increased cytotoxicity in PDOs derived from CK2 nominated tumors as compared to PDOs derived from tumors without CK2 nomination (Fig. 4D, Supplemental Fig. 6, and Supplemental Table 4). The results were further confirmed by live/dead staining (Fig. 4E). The EC_50_ in PDOs generated from tumors where KMEA nominated CK2 ranged from 88nM to 7.8μM, while PDOs generated from tumors without CK2 nomination had EC_50_ in a non-overlapping range from 10.3μM to 17.7μM, with medians of the two distributions separated by nearly a full order of magnitude (Fig. 4F).

## DISCUSSION

Motivated by the unmet need to deliver personalized cancer care to patients without druggable targets identifiable by next generation sequencing alone, we sought to apply our recent work deciphering the specificity of the human kinome [38, 56] to deconvolute patient-specific tumor phosphoproteomic data from metastatic GEP-NETs. This strategy represents an advance over existing published methodologies that use only the small subset of experimentally measured phosphopeptides matching those found in curated sets of kinase-substrate relationships derived from previous published work [34, 74]. Such curated sets are not available for most kinases, and those available likely do not contain all of the kinase’s substrates [36, 75]. Instead, by using the motif specificities of the human kinome as a handle, our KMEA approach allows for the use of all measured phosphorylation sites in the patient-derived phosphoproteomic dataset to assess the activity of nearly all human serine/threonine kinases.

Although KMEA-based analyses are employable for any tumor type, we chose to focus our initial translational efforts on GEP-NET liver metastases for several reasons. First, patients frequently succumb to liver failure secondary to parenchymal replacement by tumor [76–78]. We chose to use patient-matched uninvolved hepatic parenchyma as a control for each tumor in our analysis in order to try to identify the kinase targets least likely to impart collateral hepatoxicity, which is the most frequent single cause of safety-related drug withdrawals [79]. Second, given the currently-limited personalized therapy options [80], aggressive cytoreductive surgery is advocated by international consensus guidelines [76–78] as a means to “reset” the clock on hepatic parenchymal replacement by tumor. These procedures, as were done in the present study, provide access to large volumes of tumor that can be immediately snap-frozen in the operating room without compromising standard histopathologic examination, as would occur if primary tumors were used. Third, although recurrence following aggressive cytoreductive surgery is highly likely, most patients will not require additional therapy for one or more years after the procedure [81–86], providing a window of time for tumor evaluation and selection of patient-specific therapies. Finally, recent successes in the development of GEP-NET patient-derived organoids [87] provide an effective patient avatar on which to test KMEA predictions on GEP-NET tumor metastases.

We first elected to validate KMEA-predicted mTOR activity because it represents an ideal test case for this platform. Everolimus is an approved, highly selective allosteric inhibitor of mTORC1 that binds FKBP12 to block mTOR signaling, and it is already used in patients with advanced NETs. In the RADIANT-3 trial, everolimus improved median progression-free survival in PNETs from 4.6 to 11.0 months [62] and in RADIANT-4 it improved median progression-free survival in advanced nonfunctional gastrointestinal or lung NETs from 3.9 to 11.0 months [88]. However, everolimus is administered without a validated companion diagnostic, and approximately 20% of treated patients derive little or no benefit, underscoring the need for functional biomarkers. This discrepancy in response is the rule in oncology rather than the exception. For example, everolimus is also approved in renal cell carcinoma following progression on vascular endothelial growth factor receptor tyrosine kinase inhibitors, where it improved progression-free survival from 1.9 to 4.9 months in the RECORD-1 trial although 20% of treated patients experienced disease progression as their best overall response [89]. These observations highlight the limitations of using histology alone to infer pathway dependency and support the need for direct measurement of signaling activity for targeted therapies to supplement NGS. It is important to note the NGS matched three patients to everolimus, while KMEA identified two of these patients along with four others as having active mTOR tumor signaling.

Following our examination of mTOR, we focused on CK2 given that it was nominated in >50% of the patients included in this study. Interestingly, proteogenomic characterization of primary PNETs employed kinase-substrate enrichment analysis (KSEA) to identify CK2 activity as one of 49 kinases showing differential activation [50]. To our knowledge, this was the first report of CK2 activity in any NET, although no validation was undertaken. CK2 is a constitutively active serine/threonine kinase, with hundreds of substrates, that augments active signaling pathways, including PI3K/AKT/mTOR, IKK/NF-κB, JAK/STAT, and Wnt/B-Catenin [90], in seeming accordance with the ostensible lack of pathway drivers in this disease. Elevated CK2 expression has been identified in many epithelial solid tumors, where robust cell line and xenograft models were available to validate its elevated expression through genetic control of expression/kinase activity [91]. As compared to patient-matched uninvolved liver, we observed CK2 overexpression in nearly all GEP-NETs by IHC, but increased activity in only a fraction, highlighting the complementary nature of KMEA to standard tumor evaluations.

COT and FAM20C were among the kinase motif signatures that most strongly distinguished GEP-NET liver metastases from patient-matched uninvolved liver, and their nomination highlights both the value and the limitations of KMEA. Neither kinase currently has a clinically-approved inhibitor, but both point toward biology that is not readily captured by NGS alone. COT, encoded by MAP3K8, is a MAP kinase pathway agonist that can reactivate ERK signaling through MEK-dependent mechanisms independent of RAF. Johannessen et al. previously identified COT as a driver of resistance to RAF inhibition in BRAF-mutant melanoma, establishing it as a clinically relevant example of kinase pathway rewiring that can mediate escape from otherwise rational targeted therapy [59]. The identification of COT in our analysis suggests that MAPK pathway activation may contribute to growth or survival signaling in a subset of GEP-NET metastases. FAM20C is also particularly intriguing in this disease context because it is the major Golgi-localized kinase responsible for phosphorylation of secreted proteins [60], and NETs are defined, at least in part, by their secretory function [61]. It is likely that inhibition of FAM20C could induce significant proteotoxic stress as a novel therapeutic strategy. This illustrates the strength of KMEA as a discovery framework—identifying signaling programs that may reveal vulnerabilities exploitable through combinatorial approaches even when the nominated kinase itself is not yet directly druggable.

Though much work needs to be done to move KMEA to the “pipeline” status of NGS, we strongly believe that this is a first step that others can follow for seemingly any solid or liquid malignancy. It is important to emphasize that KMEA nominates kinase candidates of interest but does not, in isolation, establish therapeutic dependence. Kinase activity may reflect lineage state, adaptive signaling, stromal influences, or compensatory pathway activation rather than an exploitable tumor vulnerability. Accordingly, KMEA should be viewed as a prioritization framework that requires orthogonal validation. We see this approach as immediately relevant and complementary to NGS efforts, with respect to stratifying drugs to screen using patient-derived models, which are becoming a pivotal, NIH-endorsed resource for cancer care. Although not included in the present investigation, KMEA is not restricted to serine/threonine kinases. The same conceptual framework could be extended to tyrosine kinase signaling using recently developed kinase motif compendia [56], which is important given the number of available tyrosine kinase inhibitors, but the low abundance of phosphotyrosine (about 1% that of phosphoserine and phosphothreonine) and the similarity of tyrosine kinase motifs may prove limiting in this regard. In conclusion, this study demonstrates the feasibility of utilizing personalized kinase motif analysis for investigation of new kinase targets. The study carries broad implications for the use of proteomic data to help guide personalized therapies alongside standard NGS.

## MATERIALS AND METHODS

### Patient tissue acquisition

Specimens were procured under an Institutional Review Board (IRB) protocol approved by the National Institutes of Health (NIH) IRB. All patients with GEP-NET undergoing surgical procedures provided informed consent. Analyses per tumor as described below are delineated in Supplemental Fig. 1.

### TruSight Oncology 500 assay

Genomic DNA was extracted from formalin-fixed paraffin-embedded (FFPE) tissue sections after macrodissection for viable tumor content enrichment using AllPrep DNA/RNA FFPE Kit or QIAmp DNA FFPE Tissue Kit/RNeasy FFPE kit (Qiagen). 150ng total RNA from FFPE samples were used to prepare libraries with TruSeq RNA Exome kit per manufacturing instructions. Final enriched libraries were sequenced on NextSeq 550DX or NovaSeq 6000 (Illumin® Inc). Classification of pathogenicity and actionability of variants (Tier 1 to 4) was performed by QIAGEN Clinical Insight (QCI) Interpret that are based on integration of clinical and population-based bioinformatics databases, protein function prediction tools, and from experimental and clinical data reported in the biomedical literature. [92–94] A board-certified pathologist generated final report after review of the variant classification and interpretation. Limit of detection (LOD) for fusions is about 7% as determined in the analytical sensitivity testing by serially diluting 100% reference positive RNA samples with normal tissue RNA. RNA input is required at 150 ng (FFPE sample) and the targeted RNAseq reads is 60 million.

### Phosphoproteomics Analysis

Briefly, phosphoproteomics analyses of GEP-NET liver metastases and patient-matched uninvolved liver clinical specimens were performed by quantification of tandem mass tag (TMT)-labeled specimens. Details of respective experimental procedures are described below.

#### Tissue Homogenization and sample processing

A total of 20 liver metastases and 14 patient-matched uninvolved liver samples were homogenized by treatment with 1mL of lysis buffer containing EasyPep lysis buffer (Thermo Fisher Scientific #A45735), 1X Phosphatase Inhibitor (Thermo Fisher Scientific #A32957) and universal nuclease (Thermo Fisher Scientific #88700, 1µL nuclease per 500µL of buffer). The tissues were split across 2 plexes, each plex contained 10 liver metastases and 7 patient-matched uninvolved liver tissue samples. One plex was devoted to SBNET-related samples, and the other to PNET-related samples. A single PNET patient, PNET 7, had 2 samples taken from the same lesion. Assessment of mTOR, CK2A1, and CKA2 activity in these samples were extremely similar and did not differ at the 10% FDR cutoff level used in this work.Samples were homogenized using a Bead Ruptor 12 (OMNI International) with shaking at speed 4 for 20 seconds, waiting for 15 seconds, then shaking for an additional 20 seconds. Samples were quickly centrifuged at 500g for 10 seconds and the supernatant was removed from the tissue and placed in a clean 1.5mL Protein Lo-Bind Eppendorf tube. Samples were diluted to a working concentration of less than 2µg/µL.

Protein concentration was determined by the Bicinchoninic Acid (BCA) protein assay and 110µg was taken from each sample for digestion. Samples were adjusted to 200μL total volume with lysis buffer and treated with 50μL of digestion buffer containing the following composition: 60mM TCEP, 200mM chloroacetamide, and 50ng/µL trypsin/LysC in 80mM HEPES pH 8. The samples were digested at 37°C for 21hrs then a volume equal to 10µg was removed from each sample and used to create a reference pool assigned to TMTpro channel 135. Each sample and reference pool were treated with 40μL of 5µg/μL TMTpro 18-plex label (Thermo Fisher Scientific #A52045) and incubated for 1hr at 25°C. Excess TMTpro was quenched with 50μL of 5% hydroxylamine, 20% Formic acid for 10 minutes and samples within each plex were then combined and the reference pool was added to each plex. Samples were cleaned using EasyPep Maxi columns provided with the EasyPep kit (Thermo Fisher Scientific #A45734) as described in the manual. Peptides were eluted in 3mL of elution buffer and each plex was aliquoted into a 50µL aliquot (for global analysis), 4×737µL aliquots (for phospho enrichment) and dried.

Phosphopeptide enrichment was carried out sequentially using Thermo High Select TiO_2_ (#A32993) and High Select Fe-NTA (IMAC, #A32992). The 4×737µL aliquots of each plex were resuspended in the TiO_2_ Binding buffer and enriched by TiO_2_ according to the manufacturer’s protocol. The TiO_2_ elution and flowthrough were then dried and the TiO_2_ flowthrough was further enriched by IMAC according to the manufacturer’s protocol. The TiO_2_ and IMAC elutions were combined for MS analysis as described below.

#### LC/MS data acquisition

For each plex the 50µL aliquot was resuspended in 50µL 0.1% FA and the TiO2 and IMAC elutions were combined in 35µL total of 0.1% FA. For global and phosphopeptide analysis 15μL was analyzed using a Dionex U3000 RSLC in front of a Orbitrap Eclipse (Thermo Fisher Scientific) equipped with an EasySpray ion source with FAIMS Pro Duo interface. Solvent A consisted of 0.1% FA in water and Solvent B consisted of 0.1% FA in 80% ACN. Loading pump consisted of Solvent A and was operated at 7 μL/min for the first 6 minutes of the run then dropped to 2 μL/min when the valve was switched to bring the trap column (Acclaim™ PepMap™ 100 C18 HPLC Column, 3μm, 75μm I.D., 2cm, PN 164535) in-line with the analytical column (EasySpray C18 HPLC Column, 2μm, 75μm I.D., 25cm, PN ES902). The gradient pump was operated at a flow rate of 300nL/min and each run used a linear LC gradient of 5-7%B for 1min, 7- 30%B for 134min, 30-50%B for 25min, 50-95%B for 4min, holding at 95%B for 7min, then re-equilibration of analytical column at 5%B for 17min. Each sample was injected twice using two different acquisition methods that differed only in the FAIMs compensation voltages used. Both injections employed the TopSpeed method with four FAIMS compensation voltages (CVs) and a 0.75 second cycle time for each CV (3 second cycle time total) that consisted of the following: Spray voltage was 2200V and ion transfer temperature of 300 ⁰C. MS1 scans were acquired in the Orbitrap with resolution of 120,000, AGC of 4e5 ions, and max injection time of 50ms, mass range of 350-1600 m/z; MS2 scans were acquired in the Orbitrap using TurboTMT method with resolution of 15,000, AGC of 1.25e5, max injection time of 22ms, HCD energy of 38%, isolation width of 0.4Da, intensity threshold of 2.5e4 and charges 2-6 for MS2 selection. Advanced Peak Determination, Monoisotopic Precursor selection (MIPS), and EASY-IC for internal calibration were enabled and dynamic exclusion was set to a count of 1 for 15sec. The first injection used CVs of -45, -55, -65, -75 and the second used CVs of -50, -60, -70, -80.

#### Database search and post-processing analysis

For each plex the 4 MS files were batched and searched with Proteome Discoverer 2.4 using the Sequest node. Data was searched against the Uniprot Human database from August 2023 using a full tryptic digest, 2 max missed cleavages, minimum peptide length of 6 amino acids and maximum peptide length of 40 amino acids, an MS1 mass tolerance of 10 ppm, MS2 mass tolerance of 0.02 Da, variable oxidation on methionine (+15.995 Da), phosphorylation (+79.966) on serine, threonine and tyrosine, TMTpro (+304.207) on lysine and peptide N-terminus, and fixed modification of carbamidomethyl on cysteine (+57.021). Percolator was used for FDR analysis, phosphosite localization was set to 90% in the IMP-ptmRS node, and TMTpro reporter ions were quantified using the Reporter Ion Quantifier node and normalized on total peptide intensity of each channel 135). TMTpro channel assignment for conditions can be found in Supplemental Table 1.

### Kinase-Motif Enrichment Analysis (KMEA) for Phosphoproteomics

SBNET and PNET phosphoproteomic data sets were read into R (version 4.5.0) [95]. Peptides with an XCorr of 1.7 or greater were kept, and those with a lower XCorr were discarded. For each comparison of interest (either bulk SBNET or PNET liver metastases vs patient-matched uninvolved liver or an individual patient’s lesion vs. the same patient’s control) the log_2_ fold-change between the averages of the relevant samples was computed for each peptide. Phosphopeptides containing only tyrosine phosphorylations were rare and were discarded before analysis, as were phosphopeptides not detected in any of the relevant samples. The remaining phosphopeptides, each containing 1 to 3 sites of serine or threonine phosphorylation, were ranked by log fold-change, with a small proportion of positive and negative infinite changes (due to nondetection in patient matched uninvolved liver or liver metastases, respectively) recorded as one log_2_ step higher or lower than the highest non-infinite positive or negative change. Each phosphorylation site on each phosphopeptide was scored for biochemical substrate suitability using the Kinase Library (https://kinase-library.mit.edu) Sites that scored at least in the 95^th^ percentile of human phosphoserines and phosphothreonines [37] were taken to be potential kinase substrates. For each kinase, a “kinase motif set” was generated comprising all phosphopeptides with at least one potential substrate site.Kinase motif sets were analyzed using the fgsea [96] implementation of the GSEA [57] algorithm in R, with an exponent value of zero (meaning that only rank ordering of phosphopeptides’ log_2_ fold-change matters, and not the quantitative value of the change). Kinases with a significantly (FDR < 10%) positive NES in each comparison were taken to be putatively hyperactivated. In all cases, predicted CK2A1 and CK2A2 hyperactivity were concordant, and are denoted simply as CK2 throughout. KMEA-related plots were generated using the fgsea package as well as ggplot2 [97].

### Patient Derived Organoid Generation

#### Tissue Dissociation

Tumor dissociation was performed within 1-hour post-surgery. Tissues were minced finely, and necrotic/scar tissue was discarded. Minced tissue was transferred into gentleMACS C Tubes (130-093-237, Miltenyi Biotec) and dissociation was performed using the Human Tumor Dissociation Kit (130-095-929, Miltenyi Biotec) on a gentleMACS Octo Dissociator with Heaters (130-096-427, Miltenyi Biotec). After tissue was dissociated, 25 mL cold RPMI was added, and the resulting solution was filtered through Falcon cell strainers (70 µm) (352350, Corning) to remove remaining undigested pieces of tissue. ACK lysis buffer (118-156-721, Quality Biological) was used to lyse red blood cells in the cell mixture according to manufacturer’s instructions. Remaining cells were counted using a Nexcelom Cellometer Auto T4 (24107, Nexcelom) and live cell counts were used to plate organoids.

#### Organoid Culture

NET organoid culture medium was utilized for PDO development as described previously [39]. After establishing viable cell counts, 10 million viable cells/ mL were resuspended in a 50% mixture of growth factor reduced LDEV Matrigel (356231, Corning) and 50% NET organoid culture medium. Therapeutic screening of organoids was established by adding 1 µL (10,000 cells) droplets of viable tumor cells suspension to each well of a 96 well plate, followed by a 30-minute incubation at 37°C to form stable constructs. Following solidification of the gel, 200 µL of NET organoid medium was added to each well of the therapeutic screening plate. Simultaneously, 10 µL (100,000 cell) droplets were also plated into each well of a 48 well plate and cultured to observe growth and for immunohistochemistry. Following solidification of the gel, 500 µL of NET organoid medium was added to each well of the long-term culture plate Complete media changes were performed weekly. For all but one patient, model organoids were generated from the same lesion as was subjected to mass spectroscopy. For a single patient, SBNET 6, a second concurrently-resected lesion was used.

#### Drug Therapy Screening for PDOs

Following two weeks of culture, PDOs were treated with Everolimus (S1120, SelleckChem), Silmitasertib (S2248, SelleckChem), or DMSO carrier control. Drug solutions were prepared according to manufacturers’ instructions. Treatment solutions were prepared fresh before adding therapies. All the PDOs were treated with doses ranging from 0.01-50 μM for 96 hours, ending with 200 μL of media added to all wells (n≥4). Controls were treated with 0.5% DMSO. Following 96 hours of treatment, CellTiter Glo 3D assay (G9683, Promega, Madison, WI) was carried out as per manufacturer’s instructions. Further, the dose response curves were generated from the normalized values with non-linear regression curve fitting. The sigmoidal 4PL with X as (log concentration) was selected as an equation to build the dose response curves. The parameters for top, log IC_50_ and Hill slope were unconstrained, and the bottom parameter was constrained to be greater than zero (Bottom > 0). The control was added on the curve as a red dotted line and treated as a normalization factor equal to 100%. The curves were generated in GraphPad Prism version 10.6.0.

### Live and Dead staining

Live/Dead staining was performed for 1 hour with LIVE/DEAD™ Viability/Cytotoxicity Kit (Invitrogen, #L334) containing LIVE dye, 2µM Calcein-AM, and DEAD dye, 4µM ethidium homodimer with excitation wavelength of 494/528 nm and emission wavelength of 517/617 nm. The imaging was performed using an Invitrogen EVOS^TM^ FL Imaging System microscope.

### Immunohistochemistry Staining

#### SBNET and PNET liver metastases and PDO validation

Portions of the SBNET and PNET liver metastases were immediately fixed in 10% formalin (Azer Scientific, #B-PFNBF-60) after the tumor removal and blocked into formalin fixed paraffin embedded (FFPE) blocks. At day 21, matrigel containing organoids were carefully removed and placed into tubes. Organoids were washed twice with PBS and fixed in 4% paraformaldehyde for two hours. Paraformaldehyde solution was removed, and organoids were washed twice with PBS. PBS was replaced with 70% EtOH (LeicaBiosystems #3803686), following dehydration through graded alcohols, cleared in xylene, and then infiltrated with paraffin for preparing FFPE blocks. These blocks were sectioned on a microtome at 5 µM and placed onto slides. The sections were stained with Ki67(D2H10, #9027, Cell signaling, 1:300), rabbit monoclonal Synaptophysin (D8F6H, #36406, Cell signaling, 1:200) and mouse monoclonal chromogranin A (5H7, #36468, Cell signaling, 1:200) on LeicaBiosystems’ BondRXm autostainer with the following conditions: Epitope Retrieval 1 (Citrate-pH-6.0) (LeicaBiosystems #AR9961) and Epitope Retrieval 2 (EDTA; pH-8.0) (LeicaBiosystems #AR9640) for 20 minutes, and staining was carried out using the Bond Polymer Refine Detection Kit (LeicaBiosystems #DS9800). Slides were removed from the Bond autostainer, dehydrated through 70, 90, and 100% ethanol (LeicaBiosystems #3803686) for 1 minute each followed by xylene (Fischer Scientific #X3RB50) for 1 minute, and coverslipped with the xylene based CoverMount (Avantik Biogroup #SL6012-A). Further slides were scanned via NanoZoomer S60 at 40X magnification (Hamamatsu) and images were viewed using NDP.view2 software [98].

#### Casein Kinase 2 Immunohistochemistry Staining

For casein kinase 2 staining, IHC staining was performed on LeicaBiosystems’ BondRXm autostainer as described above with the following conditions : Epitope Retrieval 2 (EDTA) for 20 minutes, anti-Casein Kinase 2 alpha rabbit primary antibody (Atlas Antibodies #HPA061698, 1:25) for 60 minutes, and staining was carried out using the Bond Polymer Refine Detection Kit (LeicaBiosystems #DS9800). The Isotype antibody (Novus Biologicals #AB-105-C) was used as a negative control. Slides were scanned by NanoZoomer S60 at 40X magnification (Hamamatsu) and images were viewed using NDP.view2 software. Further, slides were reviewed and scored by a pathologist for NET morphology, tumor grading and intensity of the staining.

## Acknowledgements

This work is the result of NIH funding, in whole or in part, and is subject to the NIH Public Access Policy. Through acceptance of this federal funding, the NIH has been given a right to make the work publicly available in PubMed Central. The contributions of the NIH author(s) were made as part of their official duties as NIH federal employees, are in compliance with agency policy requirements, and are considered Works of the United States Government. However, the findings and conclusions presented in this paper are those of the author(s) and do not necessarily reflect the views of the NIH or the U.S. Department of Health and Human Services.

We gratefully acknowledge advice and comments from all members of the Yaffe laboratory during the course of this work.

## Study Approval

GEP-NET liver metastases and patient-matched uninvolved liver parenchyma clinical specimens were procured under an Institutional Review Board (IRB) protocol (000491-C and 13C-0176) approved by the National Institutes of Health (NIH) IRB. All GEP-NET patients undergoing surgical procedures provided informed consent. This study was conducted in accordance with the guidelines of the Declaration of Helsinki, Belmont report, and U.S. Common Rule.

## Author contribution

Conceptualization: JMH, MBY, BAJ, TP; Data curation: TP, BAJ, PPD, SDF, MBY; Investigation: TP, BAJ, PPD, SDF, RH, KR, KC, TL, AMJ, NR, LM, AR, ALS, ES,SS,RF, SM, KL, SX, CML, AVE, LRF, JH, MBY, JMH; Formal analysis: TP, BAJ, PPD, SDF, RH,TA, MBY, JMH; Medical and surgical patient management: JMH, JDR, JLD, AMB; Patient-derived organoid model establishment: SDF, SMS, TP; Pathology: BG and DEK; Mass spectrometry: RH and TA; Project administration: JMH and MBY; Funding acquisition: JMH and MBY; Writing – original draft: JMH, BAJ, MBY, PPD, TP, KR; Review & editing: All authors have reviewed and edited the manuscript.

## Data, Code, and Materials Availability

All KMEA code will be available through GitHub upon acceptance, and is immediately available to the reviewers upon request. The mass spectrometry proteomics data have been deposited to the MassIVE repository and are available as accession number MSV000102580.

---

**Supplemental Figure 1:**
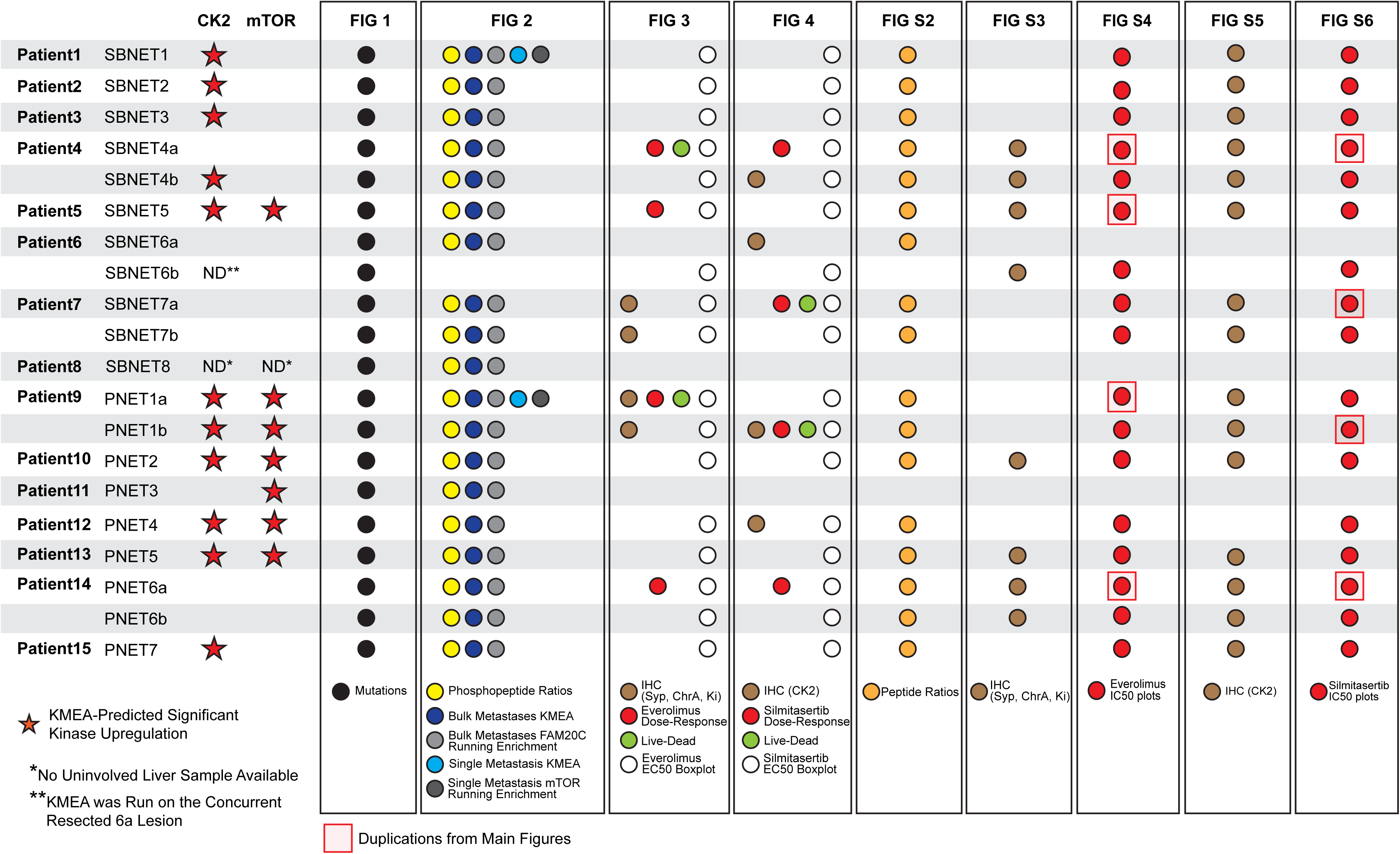
Table showing details of which patients’ SBNET and PNET liver metastases and patient-derived organoids are shown in the main and supplementary figures, as well as the predicted hyperactivation of mTOR and CK2 in each lesion.

**Supplemental Figure 2.**
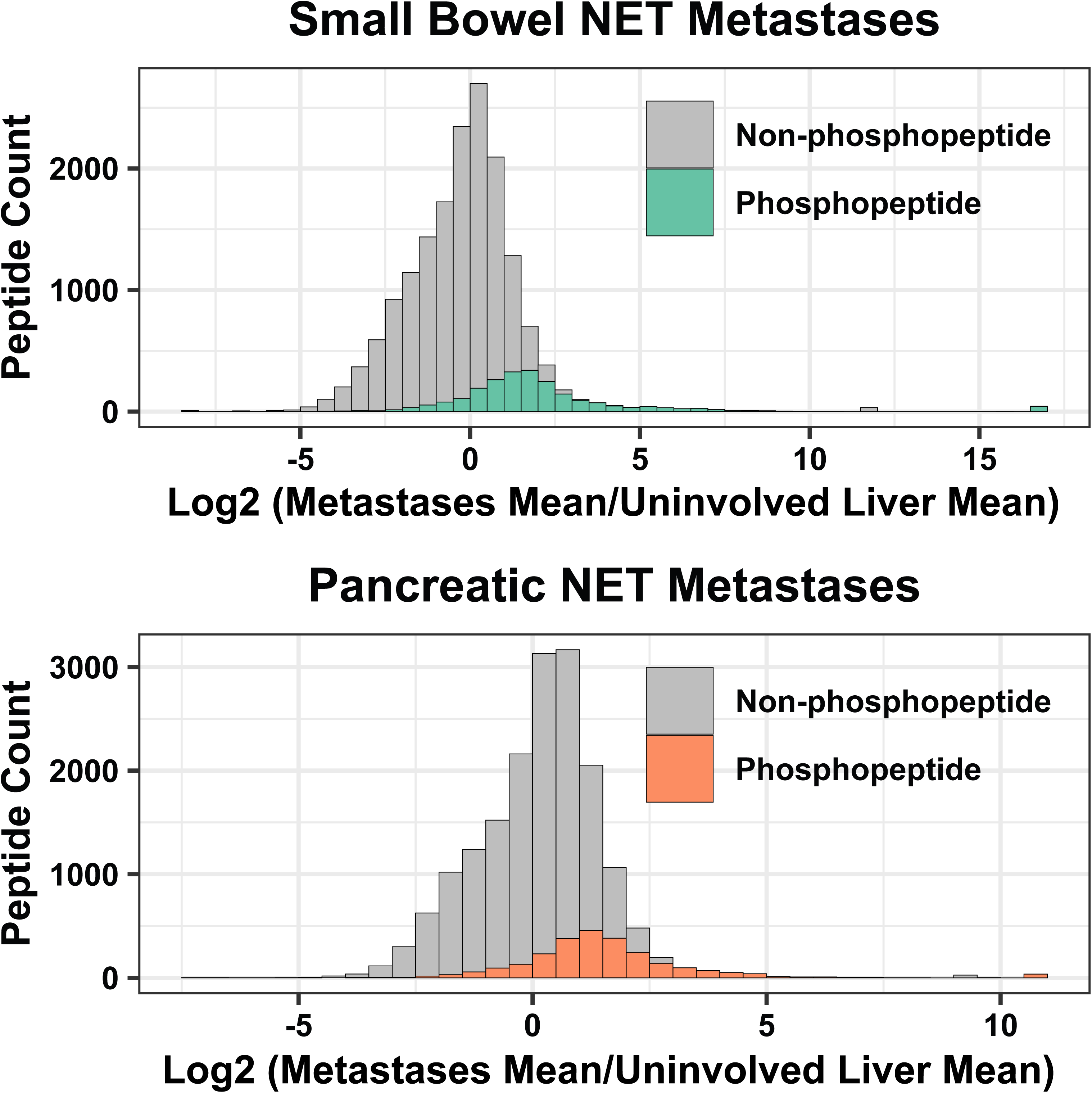
KMEA reveals potentially translatable targets in GEP-NET liver metastases. Histograms of log_2_ fold-change in phosphopeptides and non-phosphorylated peptides in the mean of liver metastases relative to the mean of patient-matched uninvolved liver as a control for bulk SBNET (top) and PNET (bottom).

**Supplemental Figure 3.**
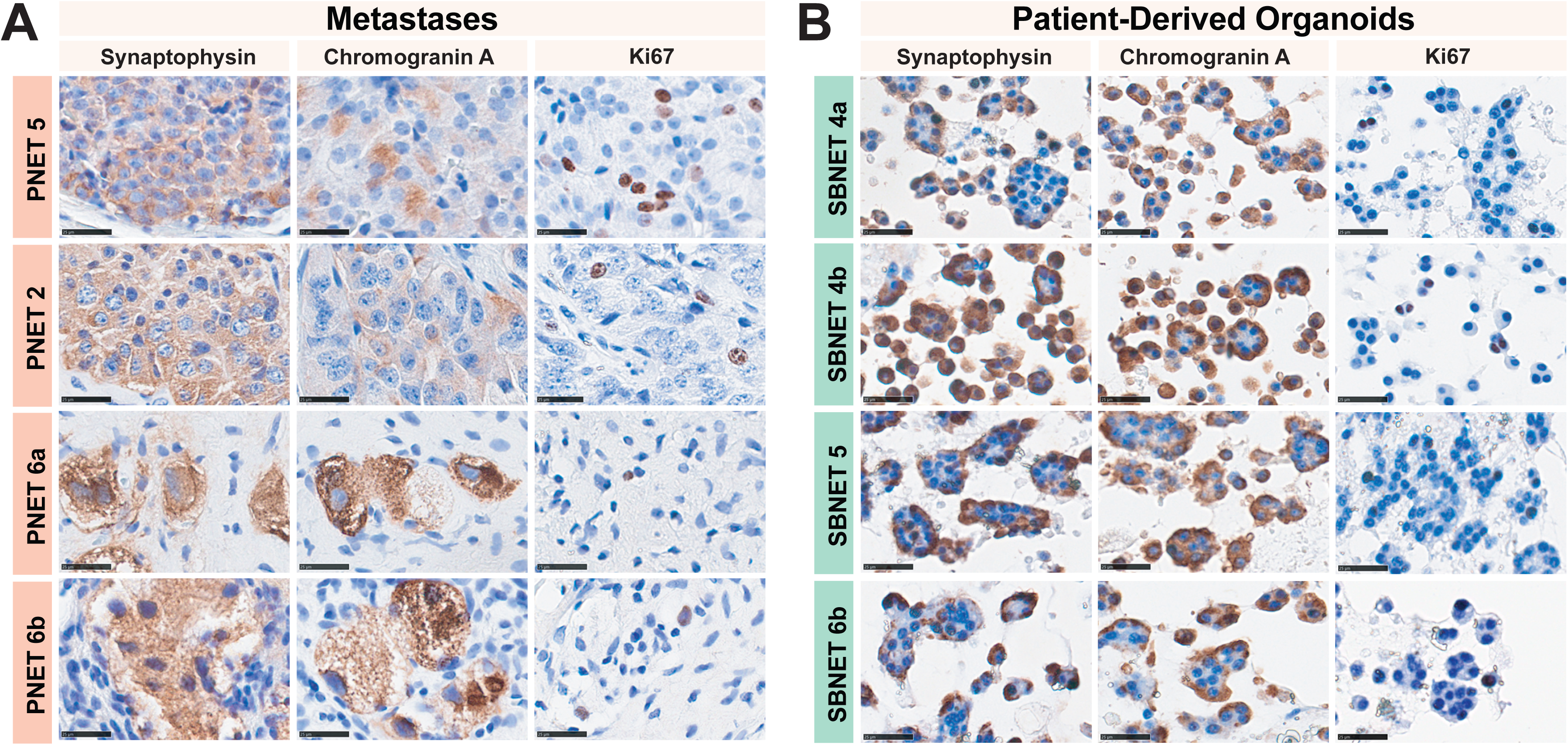
KMEA identifies mTOR therapeutic vulnerability in a subset of GEP-NET liver metastases. Representative images of Immunohistochemical characterization of synaptophysin, chromogranin A, and Ki67 in **A.** PNET patient liver metastases (*n=4*) and **B.** SBNET-PDOs (*n=4*) (right). Scale bar, 25 μm.

**Supplemental Figure 4:**
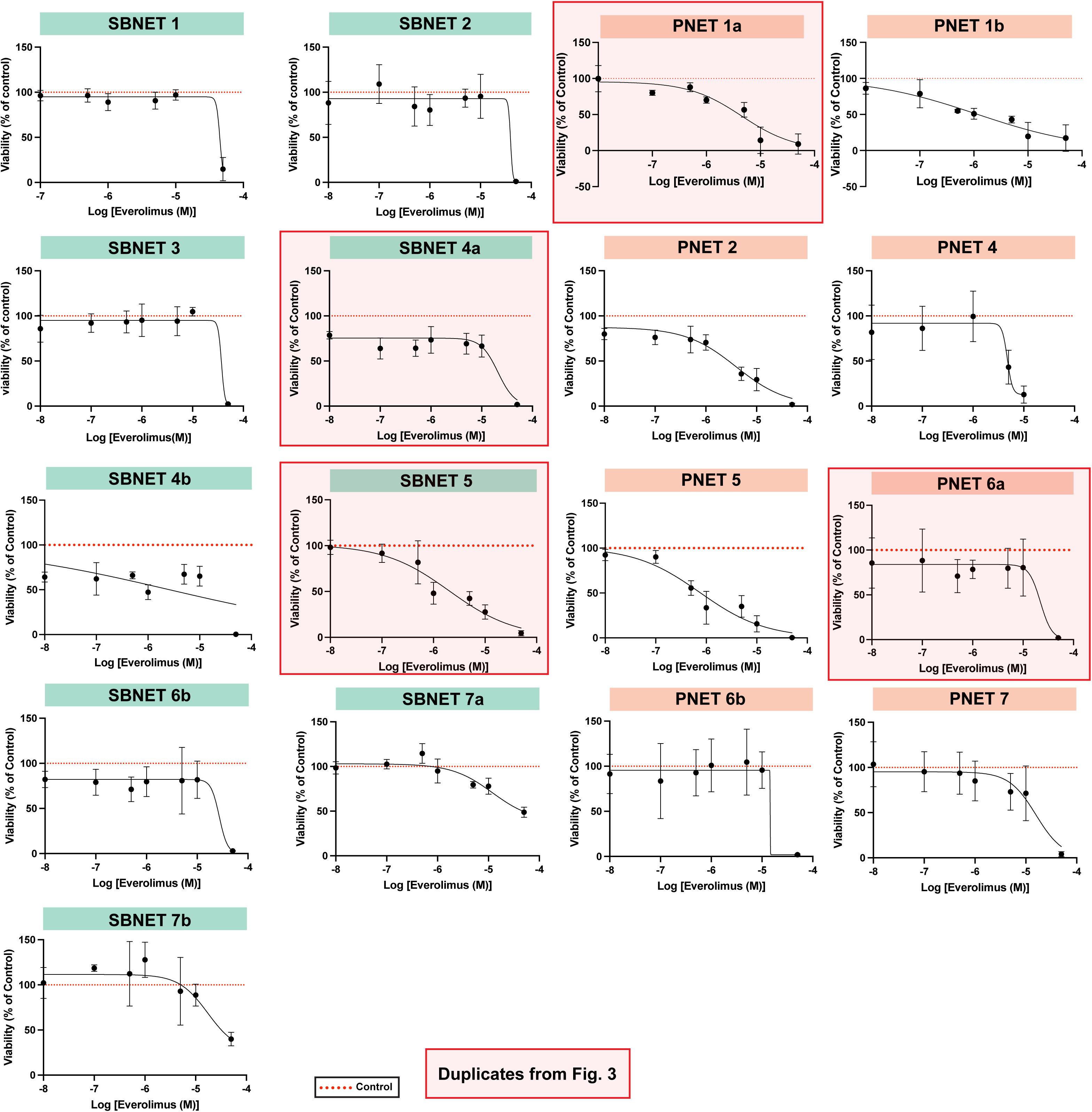
A subset of GEP-NET PDOs are everolimus-sensitive. Dose-response curves of everolimus treated SBNET (left, *n=9*) and PNET (right, *n=8*) PDOs showing percentage viability relative to control. Red dotted line represents control. Red boxed graphs are duplication from Figure 3.

**Supplemental Figure 5:**
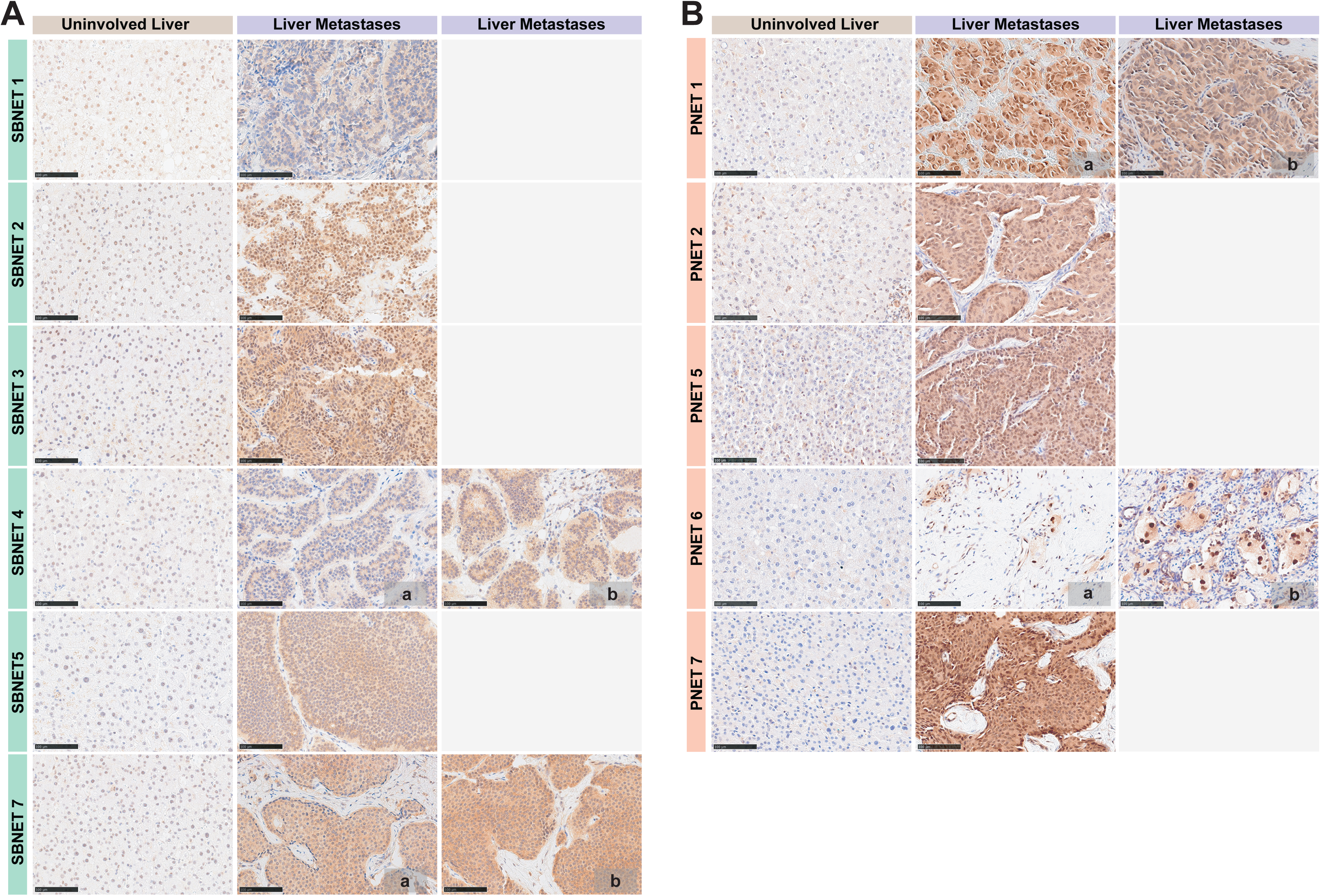
KMEA reveals casein kinase 2 as a novel therapeutic target in GEP-NET liver metastases. Representative images of Immunohistochemical characterization of Casein Kinase 2 (CK2) staining in **A**. SBNET patient-derived liver metastases (right, *n=8*) and patient-matched uninvolved liver (left, *n=6*). **B.** PNET patient-derived liver metastases (right, *n=7*) and patient-matched uninvolved liver (left, *n=5*). Where indicated, *a* and *b* denote distinct liver metastases from the same patient. Scale bar, 25 μm.

**Supplemental Figure 6:**
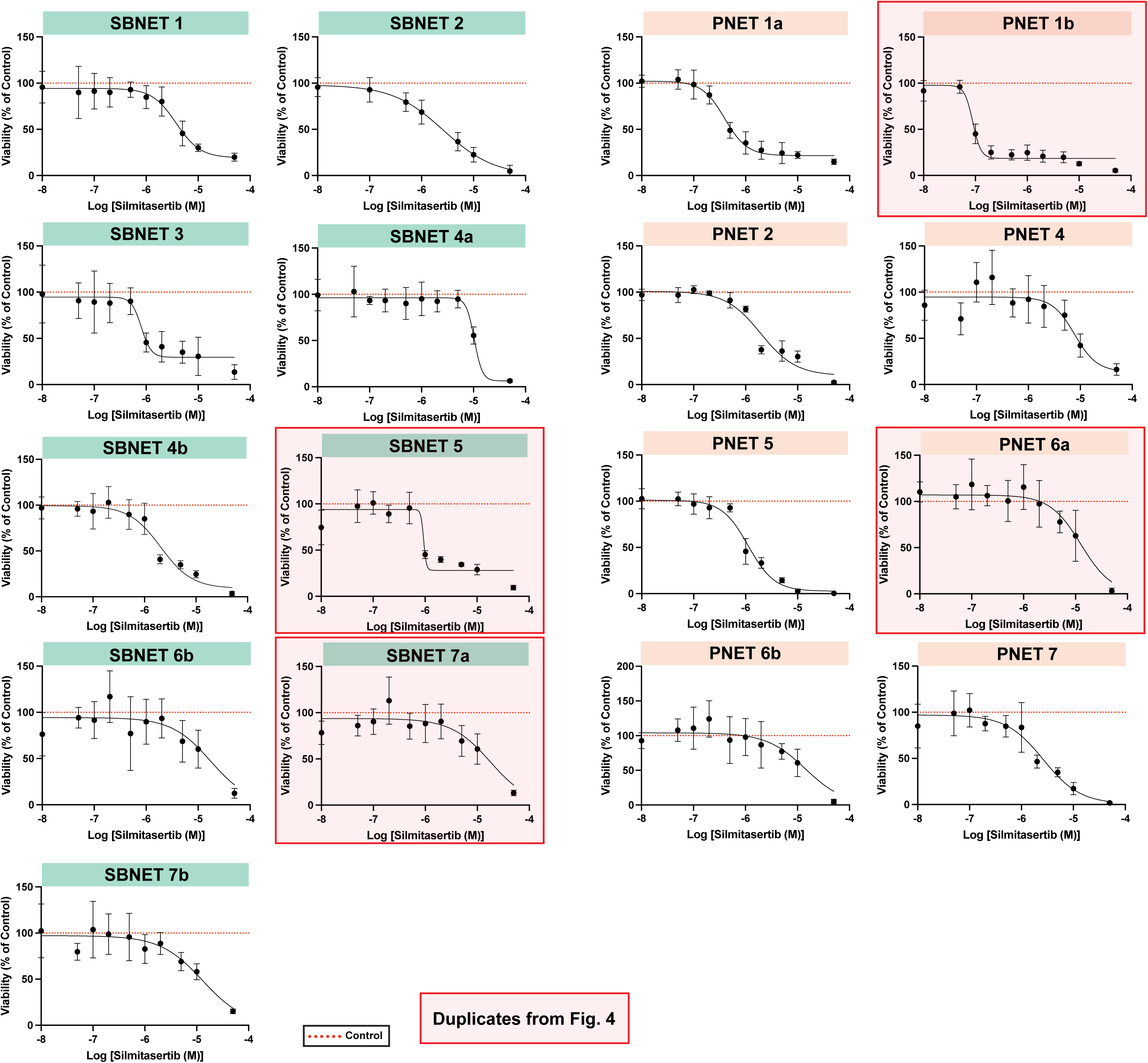
A subset of GEP-NET PDOs are silmitasertib-sensitive. Dose-response curves of silmitasertib-treated SBNET (left, *n=9*) and PNET (right, *n=8*) PDOs showing percentage viability relative to control. Red dotted line represents control. Red boxed graphs are duplication from Figure 4.

